# *E. coli* chemotaxis is information-limited

**DOI:** 10.1101/2021.02.22.432091

**Authors:** H.H. Mattingly, K. Kamino, B.B. Machta, T. Emonet

## Abstract

Organisms must acquire and use environmental information to guide their behaviors. However, it is unclear whether and how information quantitatively limits behavioral performance. Here, we relate information to behavioral performance in *Escherichia coli* chemotaxis. First, we derive a theoretical limit for the maximum achievable gradient-climbing speed given a cell’s information acquisition rate. Next, we measure cells’ gradient-climbing speeds and the rate of information acquisition by the chemotaxis pathway. We find that *E. coli* make behavioral decisions with much less than the 1 bit required to determine whether they are swimming up-gradient. However, they use this information efficiently, performing near the theoretical limit. Thus, information can limit organisms’ performance, and sensory-motor pathways may have evolved to efficiently use information from the environment.

## Main text

Organisms’ survival depends on their ability to perform behavioral tasks. These tasks can be challenging because they require that the organism measure signals in its environment and respond appropriately, in real time. Information theory is a natural language for quantifying the fidelity of measurements and responses, but it is unclear how an abstract quantity like information might limit an organism’s performance at real-world tasks. Past studies have used information theory to understand the maximum amount of information biological systems can acquire and transmit about environmental signals ^1–6^ and have shown that they can approach certain theoretical limits ^6–9^. But high information transfer is not sufficient for high performance because not all of the information contained in the signal is behaviorally relevant, and not all of it is appropriately acted on ^10^. What limits does information place on performance, and how efficiently do organisms use the information they acquire relative to these limits?

We address these questions using one of the best-understood behaviors in biology: bacterial chemotaxis. The bacterium *Escherichia coli* alternates between “runs,” which propel the cell forward, and “tumbles,” which randomly reorient its swimming direction ^11^ (Fig. 1). Along its path, *E. coli* continuously senses the concentration *c*(*t*) of chemoattractant it encounters using transmembrane receptors. Relative changes in concentration ^12^ 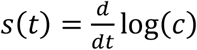 along the cell’s swimming trajectory, which we define to be the “signal”, induce changes in activity *a*(*t*) of receptor-associated CheA kinases. CheA activity goes on to modulate transitions in the cell’s motor behavior *m*(*t*) between run and tumble states via a signal transduction pathway ^13,14^. If the relative concentration of attractant is increasing (*s*(*t*) > 0), the cell tends to run for longer on average, thereby biasing its random motion up the gradient ^11^. However, noise in sensing and signal transduction corrupt the signal ^15–18^. Since the goal of chemotaxis is to climb chemical gradients, performance can be quantified by the cell’s drift velocity *v*_*d*_ up a static gradient. *E. coli* chemotaxis has been studied extensively, but the amount of information a cell acquires about chemical signals and its relationship to chemotactic performance are unclear.

**Figure 1.**
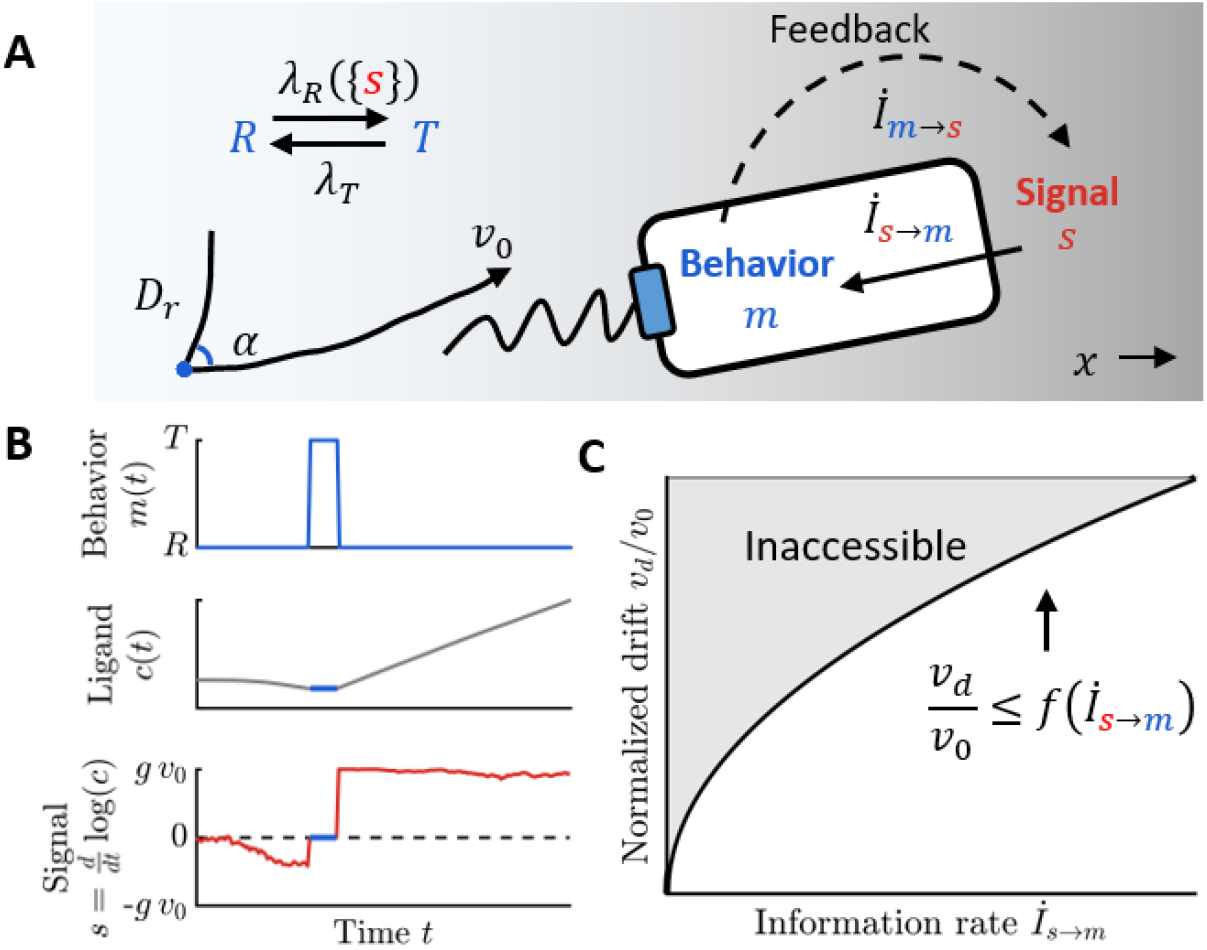
Information flow from signal to swimming behavior sets a limit on chemotaxis performance. **A)** A cell navigates chemical gradients by sensing relative changes 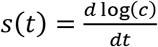 in attractant concentration *c* over time. The cell stochastically transitions between behavioral states *m*: a run state *R*, in which the cell swims with constant speed *v*_0_ and rotational diffusion *D*_*r*_; and a tumble state *T*, in which it reorients randomly with directional persistence *α* (see Fig. S1). The cell responds to past signals {*s*} by changing its transition rate *λ*_*R*_({*s*}) from run to tumble. The average tumble rate *λ*_*R*0_ and run rate *λ*_*T*_ define the fraction of time the cell spends in the run state, 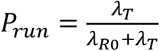. The cell’s motion creates the signal that it experiences (dashed arrow). The transfer entropy rate from behavior to signal *İ*_*m→s*_ quantifies this information flow, which is non-zero even for a non-chemotactic cell. Signal affects the cell’s behavior through *λ*_*R*_({*s*}) (solid arrow), and the transfer entropy rate from signal to behavior *İ*_*s→m*_ quantifies this information flow needed to perform chemotaxis. **B)** Time courses of behavior *m*(*t*) (top), concentration *c*(*t*) (middle), and signal *s*(*t*) (bottom) for the trajectory shown in (A). In static gradients, signals arise entirely from the cell’s motion, *s*(*t*) = *g v*_*x*_(*t*), where *g* = *d* log(*c*)/*dx* is the steepness of the log-concentration gradient and *v*_*x*_ is the cell’s up-gradient velocity. **C)** The information sent from signal to behavior *İ*_*s →m*_ sets an upper limit on chemotaxis performance, defined as the up-gradient drift speed *v*_*d*_ relative to *v*_0_. With information rate *İ*_*s →m*_, the cell’s drift speed is bounded by Eqn. 1.

Quantifying information transfer during behavioral tasks presents new challenges. First, bacteria continuously make decisions about whether to tumble. Thus, unlike most other studies in biological systems ^1–3,5,6^, we cannot use the one-shot or instantaneous mutual information ^19^ between signal *s*(*t*) and behavior *m*(*t*) as a measure of information transfer. Instead, we need an information rate ^20^. Second, the signals are generated by the cell’s own motion in the gradient ^21^. Thus, a natural extension, the mutual information rate 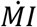 between signal *s*(*t*) and behavior *m*(*t*), is unsuitable because signal-motor correlations make it non-zero even for non-responsive cells (Fig. 1AB). We address these challenges by isolating the information that flows from signal to behavior. The mutual information rate can be decomposed into the sum of two directed information terms ^22^ (SI), the transfer entropy ^23^ rates from behavior to signal *İ*_*m→s*_ and from signal to behavior *İ*_*s →m*_, or 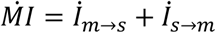. The first term *İ*_*m→s*_ quantifies the feedback of behavior onto signal. The second term, *İ*_*s →m*_, which we refer to here as the information rate, in bits/s, measures the directed influence of the signal on behavior and must be nonzero for the cell to climb the gradient.

We first used rate-distortion theory ^6,24,25^ to derive the theoretical bound on performance imposed by information transfer. To do this, we constructed a model of run-and-tumble navigation that is abstracted from *E. coli*’s molecular implementation (Fig. 1A; SI). During runs, cells swim with an effective, constant speed *v*_0_ (Fig. 1A) and lose direction with rotational diffusion coefficient *D*_*r*_. During tumbles, they reorient randomly with directional persistence *α*. In the absence of a gradient, cells switch from run to tumble and vice versa with constant rates *λ*_*R*0_ and *λ*_*T*_. In a gradient, tumbles are instead initiated at a rate *λ*_*R*_({*s*}), which depends on the past history of signals {*s*}. Throughout, we consider navigation in shallow gradients, where information is most likely to be a limitation on chemotactic performance.

Using this model, we derived how performance *v*_*d*_ and information transfer *İ*_*s→m*_ depend on the tumble rate response *λ*_*R*_({*s*}). While any response to signals implies information transfer, it does not necessarily imply high drift speed. We derived the maximum drift speed *v*_*d*_ possible given an information rate *İ*_*s →m*_ by optimizing over responses *λ*_*R*_({*s*}). In the SI, we show that in this model the optimal response only depends on the current rate of change of log-concentration, *s*(*t*). Because bacteria necessarily make comparisons of concentrations over a finite time to infer *s*(*t*) from stochastic arrivals of ligand molecules at the cell surface ^15^, they can’t implement the optimal behavioral response. Nevertheless, the performance achieved by any response is bounded by (Fig. 1C):

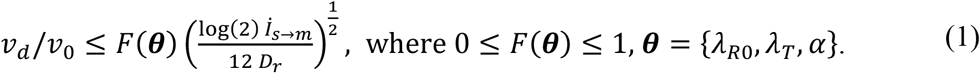

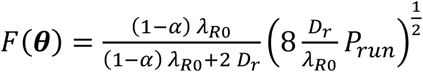 is a dimensionless function of the behavioral parameters (see Fig. 1C for a definition). This expression makes rigorous the intuition that information transfer sets a limit on how fast a cell can climb a gradient ^15,26,27^.

To quantify the bound above, we measured the rotational diffusion coefficient *D*_*r*_ and behavioral parameters θ from trajectories of swimming *E. coli* cells (Figs. S1, S2). Individual cells in a clonal population exhibit nongenetic differences in behavioral parameters θ ^28–30^, which in *E. coli* are highly correlated with *P*_*run*_, the fraction of time the cell is running *P*_*run*_ ^28,30^ (Fig. S1,S2). From this data, we find *F*(θ) = 0.531 ± 0.005 for the median phenotype (± one standard error; parameters are in Table S1). Perhaps surprisingly, the bound predicts that run-and-tumble navigation is theoretically possible with very small information rates: a hundredth of a bit per second is sufficient to climb gradients at ∼6% of the run speed. This is far less than the 1 bit per run (∼1 bit/s) required to distinguish whether concentration is currently increasing or decreasing before every tumble decision ^20^.

Our central questions are: how much sensory information do *E. coli* acquire, and how efficiently do they use that information? Eqn. 1 was derived by considering a model of *E. coli* in which signaling details have been coarse-grained away. But information about signals is acquired upstream in the *E. coli* chemotaxis pathway and must be communicated to the motors. Since this signal transduction process can only lose information ^31^, the information available at any intermediate step within the signaling pathway also places a limit on the cell’s performance, as in Eqn. 1. Intuitively, in an information-efficient cell, most sensory information acquired upstream would be preserved at the motors and contribute to gradient-climbing. If so, the cell should climb gradients at speeds approaching the theoretical bound (Fig 1C). Alternatively, cells could acquire abundant information about the signal but either lose much of it during transfer to behavior or fail to act on it appropriately. For these cells, performance would be further away from the bound. To quantify where *E. coli* lie on this spectrum, we define information efficiency, *η*, as the ratio of the cell’s chemotactic performance *v*_*d*_ to the maximum performance possible with the amount of sensory information it acquires, dictated by Eqn 1.

Quantifying *E. coli*’s information efficiency requires measuring the rate at which cells acquire information during chemotaxis. The activity *a*(*t*) of receptor-associated CheA kinases is among the first steps of *E. coli*’s signaling pathway that computes concentration changes ^14^. Therefore, we consider the rate of information transfer from signals to kinase activity *İ*_*s→a*_ ≥ *İ*_*s →m*_ to be the cell’s information acquisition rate. To quantify *İ*_*s→a*_ we measured the signal statistics cells experience during navigation, as well as the response and noise properties of the kinases in immobilized cells ^20,32^ (Fig. 2). This approach eliminates the feedback from kinase activity onto the signals cells experience, allowing us to combine these measurements to estimate the information acquisition rate *İ*_*s→a*_ during navigation (Fig. 2A; SI).

**Figure 2.**
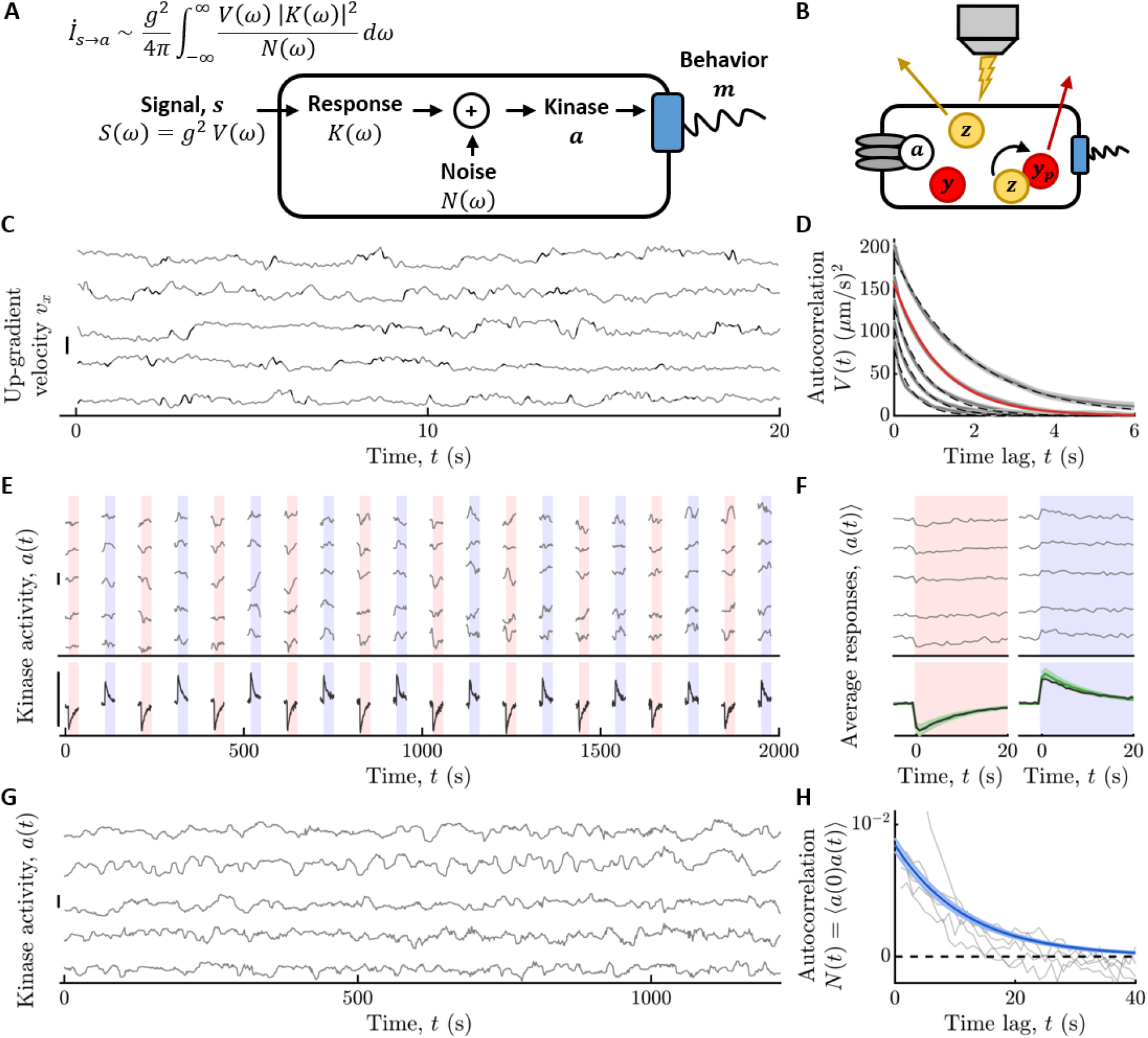
Measuring the rate of information transfer from signal to intracellular kinase. **A)** Information rate *İ*_*s →a*_ from signal *s* to kinase activity *a* depends on the signal power spectrum *S*(*ω*), the kinase frequency response *K*(*ω*), and the kinase noise power spectrum *N*(*ω*). The signal is *s*(*t*) = *g v*_*x*_(*t*), where *g* is the gradient steepness and *v*_*x*_ is the cell’s up-gradient velocity. *S*(*ω*) = *g*^2^ *V*(*ω*), where *V*(*ω*) is the power spectrum of *v*_*x*_. **B)** Kinase activity *a* was quantified from the FRET between the substrate of the kinase, CheY-mRFP, and its phosphatase, CheZ-mYFP (Methods; SI; Figs. S3-S6). **C)** (Gray) Individual cells’ *v*_*x*_(*t*) in a uniform concentration of 100 μM MeAsp. (Black: tumbles; scale bar 40 μm/s.) **D)** (Gray) Average autocorrelation of *v*_*x*_, *V*(*t*), for *P*_*run*_ = 0.93, 0.89, 0.84, 0.79, 0.74 (top to bottom; throughout, shading is ± one standard error; black dashed lines are exponential fits to *V*(*t*) = *a*_*v*_ exp(−*λ*_*tot*_ |*t*|)). (Red) Best fit to the population median bin, *P*_*run*_∼0.89 (Fig. S2; Table S1; SI). From this we computed *V*(*ω*). **E)** Immobilized cells were delivered 10 μM steps, up (red shading) and down (blue shading) (background 100 μM MeAsp). (Top, gray) Kinase activity *a*(*t*) for five cells (here and in (G), smoothed with 10^th^ order median filter, and scale bars represent Δ*a* = 0.3; Methods; SI; Fig. S5). (Bottom, black) Population average *a*(*t*) (*n* = 442 cells). **F)** Single-cell average (top, gray) and population-average (bottom, black) response functions *K*(*t*) to positive and negative stimuli. (Green) *K*(*t*) = *G* exp(−*t*/*τ*_2_) (1 − exp(−*t*/τ_1_)) *H* (*t*), where *H*(*t*) is the Heaviside step function and *G*, τ_1_, and τ_2_ are the population median parameters extracted from fits to single-cell responses. From this we computed *K*(*ω*). **G**,**H)** (Gray) Kinase activity (G) and corresponding autocorrelations (H) in single cells adapted to a constant background of 100 μM MeAsp (Methods; SI; Fig. S6). (Blue, H) 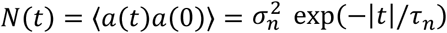, where *σ*_*n*_ and τ_*n*_ are the population median parameters extracted from fits to single-cell traces (*n* = 262 cells). From this we computed *N*(*ω*). *V*(*ω*), *K*(*ω*), and *N*(*ω*)shown in (Fig. S7).

The input signal statistics are characterized by their power spectrum *S*(*ω*). It is often difficult to know the natural signal statistics an organism experiences ^7,33^. But during bacterial chemotaxis in static gradients, the signal is generated from the cell’s own motion in the gradient. Thus, the signal power spectrum is *S*(*ω*) = *g*^2^ *V*(*ω*), where *V*(*ω*)is the power spectrum of the cell’s velocity along the gradient direction and *g* = *d* log(*c*)/*dx* is the steepness of the log-concentration gradient. Furthermore, in shallow gradients, the statistics of the cell’s motion are nearly identical to those in the absence of a gradient (SI). To quantify the *x*-velocity autocorrelation function *V*(*t*), we tracked individual swimming trajectories in a constant 100 μM background of the attractant α-methyl-aspartate (MeAsp) (Fig. 2CD; Fig. S2; Methods; SI; ∼10^4^ s of total trajectory time in the bin of median *P*_*run*_, 7 s average trajectory duration). For different values of *P*_*run*_, we fit each of the measured *V*(*t*)with decaying exponentials, 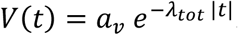. For the median phenotype, the best-fit parameters were *a*_*v*_ = 157.1 ± 0.5 (μm/s)^2^ and *λ*_*tot*_ = 0.862 ± 0.005 s^−1^. *V*(*ω*)was then computed by taking the Fourier transform of *V*(*t*) (Fig. S7).

Having quantified the input statistics, we turned to measuring the response and noise properties of CheA kinase activity (SI) using Förster resonance energy transfer (FRET) between the kinase’s substrate CheY and the phosphatase CheZ inside single cells ^17,18,34^ (Fig. 2B). In shallow gradients, cells experience weak signals, and therefore the average kinase response depends linearly on recent signals. In this regime, the response is fully characterized by how it amplifies different frequencies in the signal, *K*(*ω*), or equivalently, the response to an impulse (delta function) of signal, *K*(*t*). To measure the impulse response *K*(*t*)to MeAsp, we used a microfluidic device that allows us to rapidly switch (in ∼100 ms) the concentration of attractant delivered to hundreds of immobilized cells ^34^ (Figs. S3). To ensure cells were in the log-sensing regime ^12,35^, we first adapted them to a background of 100 μM MeAsp. We then delivered 10 positive and 10 negative 10% step changes of MeAsp concentration (corresponding to delta functions of signal *s*) (Fig. 2E; Methods), small enough that the cells’ responses were in the linear regime ^36,37^ (Fig. S5). Individual cell responses exhibited a stereotypical shape (Fig. 2F) that was well-described by a phenomenological model 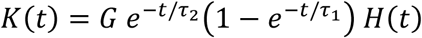, where *G* is the gain, τ_1_ is the rise time, τ_2_ is the adaptation time, and *H*(*t*) is the Heaviside step function. We fit this model to each cell’s average responses to the positive and negative stimuli simultaneously and then determined the population median values of the parameters (± one standard error; *n* = 442 cells) (Table S1; Fig. S5): *G* = 1.73 ± 0.03, τ_1_ = 0.22 ± 0.01 s, and τ_2_ = 9.9 ± 0.3 s. The value of τ_1_ that we inferred includes the CheA kinase response time, the stimulus concentration switching time, and the kinetics of CheY/CheZ binding, making it longer than the actual kinase response time. The kinase response time alone has been measured before to be τ_1_ ∼ 0.05 s ^38^, but is not directly accessible using this FRET system. After verifying that our results are not sensitive to the value of τ_1_ (SI; Fig. S8), we used the literature value in our estimate of the information rate.

We quantified the statistics of noise in kinase activity by measuring FRET in single cells in a constant background of 100 μM MeAsp (Fig. 2G; Fig. S6). These fluctuations were well-approximated by an Ornstein-Uhlenbeck process, consistent with previous measurements ^16,18^. Using Bayesian filtering (SI), we inferred the single-cell parameters of the noise model directly from the time series. These parameters determined the noise autocorrelation function 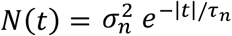 (Fig. 2H; SI), from which we computed the power spectrum *N*(*ω*) (Fig. S7). The median parameter values in the population (± one standard error; *n* = 262 cells) were *σ*_*n*_ = 0.092 ± 0.002 AU, the long-time standard deviation of the noise, and τ_*n*_ = 11.75 ± 0.04 s, the correlation time (Table S1). Note that these measurements include the effects of all noise sources upstream of the kinases, including Poisson single-molecule arrivals at the cells’ transmembrane receptors ^15^.

With the input statistics, response function, and noise, we then computed the information rate from the signal to kinase activity *İ*_*s→a*_ (Fig. 3A). Since the signal power is proportional to *g*^2^, information rate is, as well: *İ*_*s→a*_= *β g*^2^. Using our measurements above, we estimate that the *E. coli* chemotaxis system transfers information to the kinases at a rate *β* = 0.22 ± 0.03 bits/s per mm^−2^ of squared gradient steepness (SI). Thus, in shallow gradients, where concentration varies on millimeter to centimeter length scales, cells only get on the order of 10^−2^ bits/s. The bound in Eqn. 1 predicts that this should be sufficient for a run-and-tumble navigator to climb gradients at a few percent of its swimming speed. However, it is unclear how much of this information is communicated to the motors and used to navigate.

**Figure 3.**
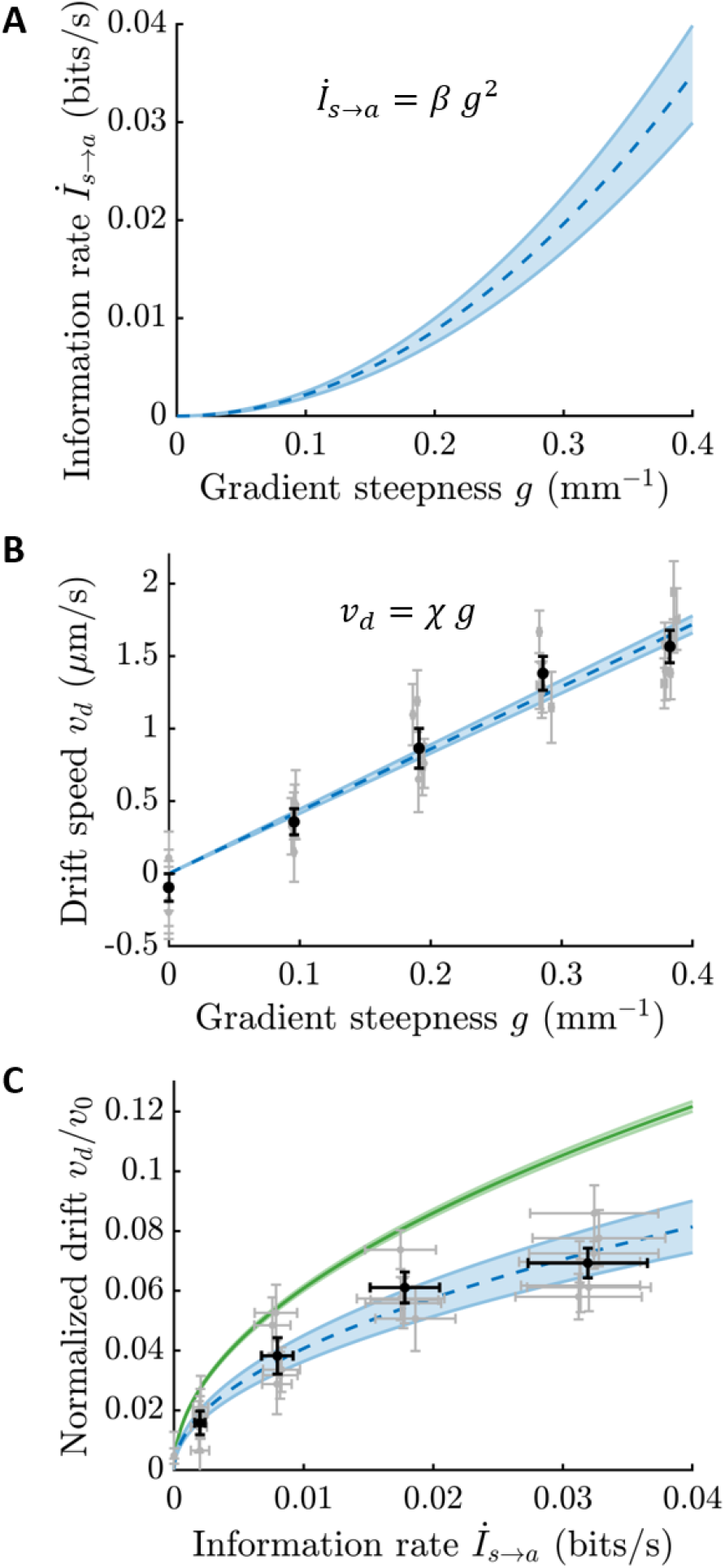
*E. coli* use information efficiently to navigate. **A)** The rate of information transfer from signal to kinase activity depends on gradient steepness *g*: *İ*_*s* →*a*_ = *β g*^2^ (*β* = 0.22 ± 0.03 bits/s / mm^−2^; Fig.2; SI). Throughout, shading and error bars indicate ± one standard error. **B)** Chemotactic performance as a function of gradient steepness *g*, in a background of 100 μM MeAsp. Gray dots: average drift speeds in individual experiments. Black dots: averages over experiments. Error bars on *g* are smaller than the markers. Population average drift speed increases linearly with gradient steepness, *v*_*d*_ = *X g* (*X* = 4300 ± 150 μm^2^/s; blue dashed line and shading; SI). **C)** From measurements of *E. coli* cells’ information rates and chemotactic drift speeds, we compared their performance to the theoretical bound (Eqn. 1). Green: predicted maximum performance given information acquisition rate *İ*_*s→a*_ (Eqn. 1). Blue: measured performance *v*_*d*_/*v*_0_ (*v*_0_ = 22.61 ± 0.07 μm/s) versus information rate *İ*_*s→a*_, obtained by eliminating *g* from the fits of *v*_*d*_(*g*) = *X g* and *İ*_*s→a*_(*g*) = *β g*^2^ to plot *v*_*d*_/*v*_0_ = *X*/*v*_0_ (*İ*_*s→a*_/*β*)^1/2^. Black and gray dots are data points from (B). Taking the ratio of the blue and green curves, we find that *E. coli* achieve drift speeds within 66 ± 5% of the theoretical limit.

To determine how efficiently *E. coli* use this information to navigate, we measured their drift speeds by tracking individual cells’ motion in gradients of varying steepness. Static, linear MeAsp gradients were constructed (Methods) in a background concentration of 100 μM, with length scales ranging from 10 mm (*g* = 0.1 mm^−1^) to 2.5 mm (*g* = 0.4 mm^−1^). From >10^5^ seconds of trajectories in each gradient condition, we estimated the average drift speed *v*_*d*_ as the time-averaged up-gradient velocity over all cells in each experiment (SI). As expected from theory in shallow gradients, the drift speeds increased linearly with gradient steepness *v*_*d*_ = *χ g*, with a proportionality constant of *χ* ∼ 4.30 ± 0.15 μm/s per mm^−1^ of gradient steepness (Fig. 3B; Fig. S9), consistent with previous measurements ^39^.

With measurements of both the information acquisition rate and the performance, we are now in a position to quantify *E. coli*’s information efficiency, *η*. For each gradient *g*, we plotted the drift speed *v*_*d*_(*g*)against the information rate *İ*_*s→a*_(*g*) (blue curve in Fig. 3C). On the same plot, we show the maximum drift speeds, given by the bound in Eqn. 1 (green curve in Fig. 3C). The ratio of these two curves is the information efficiency, 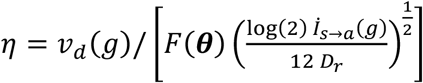. We find that *E. coli* achieve an efficiency of *η* = 0.66 ± 0.05—that is, they climb gradients at ∼66% of the maximum possible speed given the rate at which their kinases acquire information about the gradients, despite having to infer concentration changes over a finite time. The observation that *E. coli*’s efficiency is order 1 implies that much of the information about the signal contained in kinase activity is preserved in behavior and used to navigate. Thus, *E. coli* cells efficiently use information from environmental signals to perform chemotaxis, indicating that information is likely difficult for their receptor-associated kinases to acquire, limiting their performance.

Many studies of information theory in biology have focused on the maximum amount of information signaling pathways can ^1–6,40–42^. Here, we instead ask how much information an organism needs, and how much it uses, while carrying out a task, by studying information transfer in its functional context. Achieving high efficiency requires the signaling cascade to acquire, transmit, and act on information that is relevant to their behavioral task ^10^, but in many cases, which bits are relevant is not clear *a priori*. By using rate-distortion theory ^6,24,25^, we find that the bits that are relevant to bacterial chemotaxis are those that indicate how fast the attractant concentration is currently changing (see SI). This is demonstrated by our finding that maximum efficiency is achieved when the behavioral response *λ*_*R*_({*s*})is instantaneous. Furthermore, previous works measured one-shot information transfer, in bits, by biochemical networks ^1–6,40–42^. Here, we measured the rate at which *E. coli* transfer information, in bits/s, between time-varying inputs and outputs, with natural input statistics driven by the cells’ own motion. Combining this with measurements of *E. coli*’s chemotactic performance and comparing to the theoretical limit (Eqn. 1), we found that most of the information about concentration changes that flows through CheA is both relevant to and used for navigation.

The information-performance bound in Eqn. 1 depends on the cell’s swimming behavior and the physical properties of the environment in which it swims. We quantified this bound for a typical wild type cell swimming in liquid, but individual cells have different behavioral parameters **θ** ^16,28,30,29^, and the distribution of swimming phenotypes in the population depends on growth conditions. This raises the question of what is the maximum performance that can be achieved with a given information rate by any phenotype. Maximizing *F*(**θ**)in Eqn. 1 with respect to **θ** (SI), we find that the optimal agent changes direction by tumbling at the same rate as rotational diffusion ^21,43,44^ (Fig. S10). The median phenotype in our conditions tumbles more frequently than this. However, in other environments, such as semisolid agar, in which reorientations are required to escape traps ^45^, more frequent tumbling may be optimal.

*E. coli*’s proximity to the theoretical limit raises the question of how information is transmitted so reliably from continuous kinase activity into discrete transitions at the flagellar motors. Additionally, while stochastic arrival of ligand molecules at the cell’s surface ^15,46^ reduce how efficiently *E. coli* can use information by forcing them to integrate and respond to signals over a finite time, they also place an upper limit on the amount of relevant information a cell can possibly acquire. Future work will clarify what this fundamental bound on sensing is and how close *E. coli* are to it.

Information transfer is not necessarily the end-goal of biological tasks, but it is needed to perform many of them. Our results suggest organisms may be under selective pressure to efficiently use information from environmental cues to perform tasks necessary for their survival.

## Methods

### Strains and plasmids

The strain used for the FRET experiments is a derivative of *E. coli* K-12 strain RP437 (HCB33), a gift of T. Shimizu, and described in detail elsewhere ^18,34^. In brief, the FRET acceptor-donor pair (CheY-mRFP and CheZ-mYFP) is expressed in tandem from plasmid pSJAB106 ^18^ under an isopropyl β-D-thiogalactopyranoside (IPTG)-inducible promoter. The glass-adhesive mutant of FliC (FliC*) was expressed from a sodium salicylate (NaSal)-inducible pZR1 plasmid ^18^. The plasmids are transformed in VS115, a *cheY cheZ fliC* mutant of RP437 ^18^ (gift of V. Sourjik). The crosstalk coefficient for spectral bleedthrough was measured using a strain expressing CheZ-YFP from a plasmid, and that for cross-excitation was measured using a strain expressing CheY-mRFP from a plasmid, both of which are gifts from T. Shimizu. RP437, the direct parent of the FRET strain and also a gift from T. Shimizu, was used to measure behavioral parameters and chemotactic drift speeds. A mutant that can’t tumble due to an in-frame deletion of the *cheY* gene, VS100 (gift of V. Sourjik), was used to measure the rotational diffusion coefficient *D*_*r*_. All strains are available from the authors upon request.

### Cell preparation

Single-cell FRET microscopy and cell culture was carried out essentially as described previously ^18,34^. In brief, cells were picked from a frozen stock at −80°C and inoculated in 2 mL of Tryptone Broth (TB; 1% bacto tryptone, 0.5 % NaCl) and grown overnight to saturation at 30°C and shaken at 250 RPM. Cells from a saturated overnight culture were diluted 100X in 10 mL TB and grown to OD600 0.45-0.47 in the presence of 100 μg/ml ampicillin, 34 μg/ml chloramphenicol, 50 μM IPTG and 3 μM NaSal, at 33.5°C and 250 RPM shaking. Cells were collected by centrifugation (5 min at 5000 rpm, or 4080 RCF) and washed twice with motility buffer (10 mM KPO4, 0.1 mM EDTA, 1 μM methionine, 10 mM lactic acid, pH 7), and then were resuspended in 2 mL motility buffer. Cells were left at 22°C for 90 minutes before loading into the microfluidic device. All experiments, FRET and swimming, were performed at 22-23°C.

For swimming and chemotaxis experiments, cells were prepared identically. Saturated overnight cultures were diluted 100X in 5 mL of TB. After growing to OD600 0.45-0.47, 1 mL of cell suspension was washed twice in motility buffer with 0.05% w/v of polyvinylpyrrolidone (MW 40 kDa) (PVP-40) added. Washes were done by centrifuging the suspension in an Eppendorf tube at 1700 RCF (4000 RPM in this centrifuge) for 3 minutes. After the last wash, cells were resuspended with varying concentrations of MeAsp (see below).

### Microfluidic device fabrication and loading for FRET measurements

Microfluidic devices for the FRET experiments ^34^ were constructed from polydimethylsiloxane (PDMS) on 24 x 60 mm cover glasses (#1.5) following standard soft lithography protocols ^47^. Briefly, the master molds for the device were created with a negative SU-8 photoresist on 100-mm silicon wafers. Approximately 16-μm-high master molds were created. To fabricate the device, the master molds were coated with a 5-mm-thick layer of degassed 10:1 PDMS:curing agent mixture (Sylgard 184, Dow Corning). The PDMS layer was cured at 80 °C for 1 hour, and then cut and separated from the wafer, and holes were punched for the inlets and outlet. The punched PDMS layer was further cured at 80 °C for > 2 hours. Then, the PDMS was cleaned with transparent adhesive tape (Magic Tape; Scotch) followed by rinsing with (in order) isopropanol, methanol, and Millipore-filtered water. The glass was rinsed with (in order) acetone, isopropanol, methanol, and Millipore-filtered water. The PDMS device was tape-cleaned an additional time before the surfaces of the device and coverslip were treated in a plasma bonding oven (Harrick Plasma). After 1 min of exposure to plasma under vacuum, the device was laminated to the coverslip and then baked at 80°C hotplate for > 30 min to establish a covalent bond.

Sample preparation in the microfluidic device was conducted as follows. Five inlets of the device (Fig. S3) were connected to reservoirs (Liquid chromatography columns, C3669; Sigma Aldrich) filled with motility buffer containing various concentrations of α-methyl-aspartate (MeAsp) through polyethylene tubing (Polythene Tubing, 0.58 mm id, 0.96 mm od; BD Intermedic). The tubing was connected to the PMDS device through stainless steel pins that were directly plugged into the inlets or outlet of the device (New England Tubing). Cells washed and suspended in motility buffer were loaded into the device from the outlet and allowed to attached to the cover glass surface via their sticky flagella by reducing the flow speed inside the chamber. The pressure applied to the inlet solution reservoirs was controlled by computer-controlled solenoid valves (MH1; Festo), which rapidly switched between atmospheric pressure and higher pressure (1.0 kPa) using a source of pressurized air. Only one experiment was conducted per device.

### Single-cell FRET imaging system

FRET imaging in the microfluidic device was performed using an inverted microscope (Eclipse Ti-E; Nikon) equipped with an oil-immersion objective lens (CFI Apo TIRF 60X Oil; Nikon). YFP was illuminated by an LED illumination system (SOLA SE, Lumencor) through an excitation bandpass filter (FF01-500/24-25; Semrock) and a dichroic mirror (F01-542/27-25F; Semrock). The fluorescence emission was led into an emission image splitter (OptoSplit II; Cairn) and further split into donor and acceptor channels by a second dichroic mirror (FF580-FDi01-25×36; Semrock). The emission was then collected through emission bandpass filters (FF520-Di02-25×36 and FF593-Di03-25×36; Semrock) by a sCMOS camera (ORCA-Flash4.0 V2; Hamamatsu). RFP was illuminated in the same way as YFP except that an excitation bandpass filter (FF01-575/05-25; Semrock) and a dichroic mirror (FF593-Di03-25×36; Semorock) were used. An additional excitation filter (59026x; Chroma) was used in front of the excitation filters. To synchronize image acquisition and the delivery of stimulus solutions, a custom-made MATLAB program controlled both the imaging system (through the API provided by Micro-Manager ^48^) and the states of the solenoid valves.

### Procedure for measuring the linear response functions

All experiments were performed in a background MeAsp concentration of *c*_0_ = 100 μM. Measurements were made in single cells. First, the FRET level at minimum kinase activity was measured by delivering a saturating stimulus (1 mM MeAsp plus 100 µM serine ^49^) for 10 seconds. Immediately afterwards, the FRET level at maximum kinase activity was measured by delivering motility buffer with no attractant (0 µM MeAsp, 0 µM serine) for 5 seconds. When cells are adapted to 100 µM MeAsp, removing all attractant is sufficient to elicit a maximal response ^18,36^. Donor excitation interval (i.e., measurements of *I*_*DD*_ and *I*_*DA*_; see SI) was 0.5 seconds and acceptor excitations (i.e., measurements of *I*_*AA*_; See SI) were done before and after the set of donor excitations. After this, the concentration of MeAsp was returned to the background *c*_0_, and no serine was delivered to the cells for the rest of the experiment. Imaging was then stopped and cells were allowed to adapt to the background for 120 seconds.

After this, a series of stimuli were delivered to the cells in the microfluidic device (see Figure 2E for stimulus protocol). Importantly, the cells were only illuminated and imaged for part of the experiment in order to limit photobleaching. First, cells were imaged for 7.5 seconds in the background concentration *c*_0_. Then, the concentration of MeAsp was shifted up to *c*_+_ = 110 μM for 30 seconds and imaging continued. Donor excitation interval was 0.75 seconds and acceptor excitations were done before and after the set of donor excitations. After this time, imaging was stopped and the MeAsp concentration returned to *c*_0_ for >60 seconds to allow cells to adapt. Then, the same process was repeated, but this time shifting MeAsp concentration down to *c*_−_ = 90 μM. Alternating up and down stimuli were repeated 10 times each.

FRET levels at minimum and maximum kinase activity were measured again at the end of the experiment. The whole imaging protocol lasted <2200 seconds. In total, cells spent <60 minutes in the device, from loading to the end of imaging. Analyses of these data are described in the SI.

### Procedure for measuring the noise statistics

Spontaneous fluctuations in kinase activity were also measured in a background MeAsp concentration of *c*_0_ = 100 μM. Measurements were made in single cells. FRET levels at minimum and maximum kinase activity were measured at the beginning and the end of each experiment, as described above. As above, after these measurements, imaging was then stopped and cells were allowed to adapt to the background for 120 seconds. After this, cells were imaged for about 1200 seconds. Throughout, donor excitations (i.e., measurements of *I*_*DD*_ and *I*_*DA*_; see SI) were done every 1.0 second, except when it was interrupted by acceptor excitations (i.e., measurements of *I*_*AA*_; see SI), which were conducted every 100 donor excitations. The whole imaging protocol lasted <1400 seconds. In total, cells spent about < 60 minutes in the device, from loading to the end of imaging. Analyses of these data are described in the SI.

### Procedure to measure swimming and behavioral parameters

After the second wash, cells were centrifuged again and resuspended in motility buffer containing 100 µM MeAsp. Then, the cell suspension was diluted to an OD600 of 0.00025. The cell suspension was then loaded into µ-Slide Chemotaxis devices (ibidi; Martinsried, Germany), the same type of device used to create static gradients, described below. However, instead of tracking cells in the gradient region, we tracked their swimming in one of the large reservoirs, which are roughly 750 µm deep. 1000-s movies of swimming cells were recorded on a Nikon Ti-E Inverted Microscope using a CFI Plan Fluor 4X objective (NA 0.13). This objective’s depth of field is about ±18 µm, much shorter than the depth of the chamber. Adjusting the focal plane to the middle of the chamber made cells that were swimming near the ceiling or floor of the device, which could experience hydrodynamic interactions that affect their behavior ^50^, not visible in the movie. At the same time, this lower magnification objective allowed us to collect relatively longer swimming trajectories. Movies were captured around 30 minutes after loading cells into the chamber to mimic the gradient experiments below. Images here and below were captured using a sCMOS camera (ORCA-Flash4.0 V2; Hamamatsu). Analyses of these data are described in the SI. Five biological replicates were done for behavioral parameter measurements, and four biological replicates were done for measuring *D*_*r*_.

### Procedure to measure chemotactic drift speeds

Chemotaxis experiments were performed in µ-Slide Chemotaxis devices (ibidi; Martinsried, Germany). These devices generate a linear gradient between two concentration reservoirs that is stable for a long time. After the second wash, the cell suspension was split into two Eppendorf tubes, 0.5 mL each. After one more centrifugation, one tube of cells was resuspended in 1 mL of motility buffer with 100 µM of MeAsp, to be injected into the “low-concentration reservoir”, and the other was resuspended in 1 mL of motility buffer with 2 µM of fluorescein and varying concentrations of attractant, to be injected into the “high-concentration reservoir”. Cells in both tubes were diluted to OD 0.001 for each experiment. Loading cells in both reservoirs ensured that the concentration of cells throughout the experimental device was approximately uniform. This limited the effects of potential biases that could arise from observing a finite field of view.

Using a background concentration of at least 100 µM MeAsp ensured that the cells were in the log-sensing regime ^12^. The “high” concentrations of MeAsp used were 110.5 µM, 122.1 µM, 135.0 µM, and 149.2 µM. With 1 mm separating the two reservoirs, these concentrations produced linear gradients that approximated shallow exponentials gradients with steepness of roughly: *g* = {0.1, 0.2, 0.3, 0.4} mm^−1^. *g* was calculated from 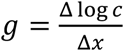, where Δ log *c* is the difference in log concentrations between the two reservoirs, and Δ*x* is the distance between them. This is exactly the average steepness of log-concentration across with the width of the channel. To see this, the steady state concentration profile is linear, 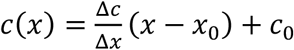, where Δ*c* is the difference in concentration between the two reservoirs, *x*_0_ is the midpoint between them, and *c*_0_ is the concentration at *x* = *x*_0_. From this, the gradient of log concentration depends on position *x* and can be computed from 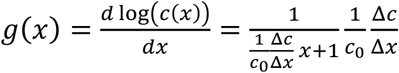, where we have defined a reference frame where *x*_0_ = 0. Averaging over the channel by integrating over *x* from −Δ*x*/2 to Δ*x*/2 and dividing by 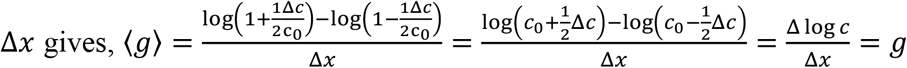. Close to the low-concentration reservoir, *g*(*x*) is larger than *g*, and vice versa near the high-concentration reservoir, but these errors are small and approximately cancel each other out when we average drift speeds of cells across the channel.

To load the device, first the reservoirs were sealed with the manufacturer’s tabs. Cell suspension with 100 µM MeAsp was injected into the channel where the gradient would form. Excess liquid in the inlets was removed. Then one tab from each reservoir was removed, and the gradient channel was sealed with tabs. The left reservoir was then fully unsealed, and the right reservoir was sealed with tabs. 60-65 µL of cell suspension with 100 µM MeAsp was injected into the left reservoir, and then both inlets of that reservoir were sealed with tape or tabs. Care was taken to make sure there were no bubbles in reservoir at the inlets. Then, the right reservoir was unsealed, and 60-65 µL of cell suspension with the higher concentration of MeAsp was injected. A timer was then immediately started. The right reservoir was then sealed.

Cells were imaged by phase contrast with a CFI Plan Fluor 10X objective (NA 0.30). The depth of the gradient region of the device is 70 µm, and the depth of field of the objective is about ±4 µm. Focusing on the middle of the chamber with this objective filtered out cells that could be interacting with the ceiling or floor surfaces. Images of fluorescein were taken every 5 minutes using a CFI Plan Fluor 4X objective (NA 0.13) through a YFP filter cube (Chroma 49003), illuminated by a LED (SOLA SE, Lumencor) with an exposure time of 100 ms. Since the diffusivity of fluorescein is similar to (slightly lower than) that of MeAsp (MW of fluorescein is 376 kDa; MW of MeAsp is 147 kDa; Sigma Aldrich), we used fluorescein as an indication of when the attractant gradient was stable and linear in the observation region between the two reservoirs, as has been done before ^51,52^. Once the fluorescein profile was stable for several time points (typically around 50-60 minutes after loading), a 1000-second phase contrast movie was recorded at 20 FPS using the 10X phase contrast objective. Before the recording, the transmitted light illumination was adjusted to minimize the number of saturated pixels. After the recording, an additional image of the fluorescein profile was recorded, and the cells were observed to check that they were still swimming normally. Analyses of these data are described below. At least five biological replicates were performed for each gradient steepness.

## Acknowledgements

We thank Jeremy Moore and Xiaowei Zhang for help setting up the experimental assays. We also thank Katja Taute and Marianne Grognot for helpful discussions about the gradient experiments. We thank Pieter Rein ten Wolde for providing detailed feedback on an earlier version of this manuscript, as well as Artur Wachtel, Isabella Graf, and Damon Clark for providing comments. We acknowledge Tom Shimizu and Victor Sourjik for bacteria strains and Rafael Gomez-Sjoberg, Microfluidics Lab, for providing information and software to control the solenoid valves in the microfluidic setup.

## Contributions

HM and KK contributed equally to this work. HM, KK, BM and TE designed the research. HM and BM derived the theoretical bound with inputs from TE and KK. HM performed the experiments tracking bacteria. KK performed the single cell FRET experiments. HM, KK, and TE validated the data. HM, KK, BM, and TE discussed the data analysis. HM and KK performed the data analysis. HH, KK, BM and TE wrote the initial draft and all revisions.

